# Dynamic multimodal interactions in navigating wood ants: What do path details tell us about cue integration?

**DOI:** 10.1101/2019.12.31.891689

**Authors:** Cornelia Buehlmann, Alexia Aussel, Paul Graham

## Abstract

Ants are expert navigators using multiple cues from multiple sensory modalities to navigate successfully. Here, we present the results of systematic studies of multimodal cue use in navigating wood ants, *Formica rufa*. Ants learnt to navigate to a feeder that was defined by an olfactory cue, visual cue and airflow presented together. When ants learnt to find a feeder that was placed in the centre of the visual cue, well-trained ants were not anymore able to accurately approach the feeder when either the olfactory or visual cue were removed in tests. This confirms that some form of cue binding has taken place. However, in a visually simpler task with the feeder located at the edge of the visual cue, ants still approached the feeder accurately when individual cue components were removed. Hence, cue binding is flexible and depends on the navigational context. In general, cues act additively in determining the ants’ paths accuracy, i.e. the use of multiple cues increased navigation performance. Moreover, across different training conditions, we saw different motor patterns in response to different sensory cues. For instance, ants had more sinuous paths with more turns when they followed an odour plume but did not have any visual cues. Having visual information with the odour enhanced performance and therefore positively impacted on plume following. Interestingly, paths characteristics of ants from the two multimodal groups with a different visual task were different, suggesting that the observed flexibility in cue binding may be a result of the ants’ movement characteristics.

**Summary statement:** We investigated the impact of multimodal information on navigating ants. Ants showed flexible response to multimodal information depending on the sensori-motor contingencies of the navigation task.

## Introduction

Ants are remarkable navigators with their efficiency coming from the coordinated implementation of a set of navigational strategies (Knaden & Graham 2016). A common strategy seen in many ant species is to utilise their social nature and develop pheromone trail networks between the nest and reliable food locations (Czaczkes *et al.* 2015). In addition, or instead of trail pheromones in some ant species, individual ants are equipped with personal navigational strategies (Wehner 2003; Collett *et al.* 2013; Knaden & Graham 2016). Such solitarily foraging ants rely on path integration (PI) and learnt information from the environment. PI is an innate behaviour that allows individuals to explore unfamiliar terrain while being safely connected to their nest (Heinze *et al.* 2018; Collett 2019). As ants become familiar with an environment, they also learn and use visual and olfactory information to guide routes (visual: (Collett *et al.* 1992; Kohler & Wehner 2005; Graham & Collett 2006; Wystrach *et al.* 2011; Mangan & Webb 2012), olfactory: (Buehlmann *et al.* 2015)) and drive searches for the nest (visual: (Wehner & Räber 1979; Wehner *et al.* 1996; Narendra *et al.* 2007), olfactory: (Steck 2012)) or a familiar feeder (visual: (Durier *et al.* 2003; Collett *et al.* 2014; Buehlmann *et al.* 2016), olfactory: (Huber & Knaden 2018)). We have a relatively good understanding of the behaviour of ants when undertaking visual navigation (Zeil 2012; Collett *et al.* 2013; Graham & Philippides 2017). However, we have only a few behavioural descriptions of olfactory navigation as a personal navigation strategy rather than in a social, pheromone trail, context (Steck *et al.* 2009; Buehlmann *et al.* 2015; Huber & Knaden 2018), and we know even less about ant behaviour during multimodal cue use during navigation.

Multiple navigational strategies often provide redundant information in experienced ants, e.g. path integration and visual guidance can influence behaviour simultaneously (Narendra 2007; Reid *et al.* 2011; Collett 2012; Legge *et al.* 2014), resulting in intermediate headings when directional cues are experimentally set in conflict (reviewed in (Wehner *et al.* 2016)). This means that navigational information is processed by separate guidance systems with convergence at the level of the behavioural output (Cruse & Wehner 2011; Hoinville & Wehner 2018). Furthermore, detailed path analyses from another set of experiments have shown that PI also interacts with other modalities by generating different movements for home vectors of different lengths, and mediating an indirect interaction between PI and the potential for learning and using visual or olfactory cues (Buehlmann *et al.* 2018).

Another way of studying multimodal navigation is looking at learning rates for ants that are trained to find an important location as defined by multimodal cues. In one such experiment, *Cataglyphis* desert ants were trained to locate their nest using either visual cues, olfactory cues or both visual and olfactory cues together (Steck *et al.* 2011). One result was that these ants learnt bimodal cues much faster than a single cue, with another finding being that bimodal landmarks were first learnt as their individual components but later stored as a holistic unit. Hence, although initially the presence of a second sensory cue enhances the learning performance of a unimodal cue, the components of the bimodal cue are fused together after several training trials and ants will no longer respond to either of the components presented alone (Steck *et al.* 2011).

At the coarse scale, behavioural outputs of multimodal interactions are adaptive and produce ecologically relevant behaviour (e.g. (Buehlmann *et al.* 2011; Cheng *et al.* 2012)). We also know that at a fine motor scale, different sensory modalities require different patterns of movement (Graham & Collett 2002; Lent *et al.* 2010; Buehlmann *et al.* 2014; Wystrach *et al.* 2014). In recent years we have gained a good understanding of visual navigational mechanisms by studying path details of wood ants in the lab (Judd & Collett 1998; Graham & Collett 2002; Lent *et al.* 2010; Lent *et al.* 2013; Buehlmann *et al.* 2016). Using that well-established system, we investigate here the details of multimodal interactions by studying the paths of wood ants guided by both multimodal and unimodal cues. Firstly, we ask if navigation is improved when using multimodal cues. Secondly, we ask if ants use different or more complex movement patterns in the presence of multimodal cues. Thirdly, we ask if the cue binding previously described (Steck *et al.* 2011) can be explained by the ants’ path characteristics and under what circumstances some behavioural flexibility is retained.

## Methods

### Ants

Experiments were performed with laboratory kept wood ants *Formica rufa* collected in Broadstone Warren, East Sussex, UK. Ants were kept in the laboratory under a 12 h light: 12 h darkness cycle and constant temperature of 25-27°C. They were fed ad libitum with sucrose and dead crickets. During the experiments, food was limited to a minimum to increase their foraging motivation, but they had access to water all the time.

### Experimental setup

General experimental procedures followed those described previously (Buehlmann *et al.* 2016). Individually marked ant foragers were taken from the nest and released in the centre of a circular platform (120 cm in diameter) that was surrounded by a cylinder (diameter 3 m, height 1.8 m) with white walls. Ants learnt to find a drop of sucrose on a microscope slide that was located 45 cm away from the centre. The feeder location was defined by multiple cues which varied between the different experiments (Fig. 1). The following cues were used: visual cue (V, black rectangle placed on the edge of the platform, 60 cm wide and 30 cm high, 60° wide and 31° high from platform centre), airflow (A, constant air flow produced by an Tetra APS 50 aquarium pump, connected to a 62 cm long silicon tube (d_outer_: 7 mm, d_inner_: 5 mm) with its end placed in the platform at r = 50 cm and pointing towards the centre) and an odour not innately attractive to the ants (O, drop of miaroma 100% pure essential pine oil pipetted on a small piece of filter paper (∼ 1 cm × 1 cm) placed 1 cm in front of the feeder and renewed every 30 mins). For a visualisation of the odour plume see Fig. 1D. These cues were presented individually (V and A) or in combinations (VA, OA and VOA). In the VOA condition, the food was either placed in the centre (VOA_Centre_) or at the left edge (VOA_Edge_) of the visual cue. Each group of ants was trained to only one of the 6 different experimental conditions (VOA_Centre_, VOA_Edge_, VA, OA, V or A). To avoid ants relying on cues other than V, O or A, the cues and the feeder were rotated together to a new position within a 120° sector after the end of each training round. Ants performed approximately 10 group trainings before being trained individually. For individual training, ants were put separately into a 6.5 cm diameter, cylindrical holding chamber in the centre of the platform. The ant was released from the holding chamber by remotely lowering the wall. Once the ant had reached the sucrose slide and started to feed, the ant was transferred into a feeding box and the next ant was released. The ants were recorded using a tracking video camera (Trackit, SciTrackS GmbH) which provided the ant’s position on the platform and the orientation of its body axis every 20 ms for analysis. All individual training runs were recorded. After approximately 14 training rounds, ants approached the feeder quite directly and once this moment was reached, tests were introduced in the VOA experiments after every 3 or 4 training trials. In these tests, either the olfactory (O) or visual (V) cue was removed from the multimodal cue combination (O removed in VOA, V+O-A+; V removed in VOA, V-O+A+) and the ants’ paths were recorded. Furthermore, conflict tests were introduced in some of the experiments. Here, either OA (for VOA_Centre_ ants) or A (for VA ants) was displaced 20° to the left of its familiar position and the ants were recorded as described above.

**Figure 1:**
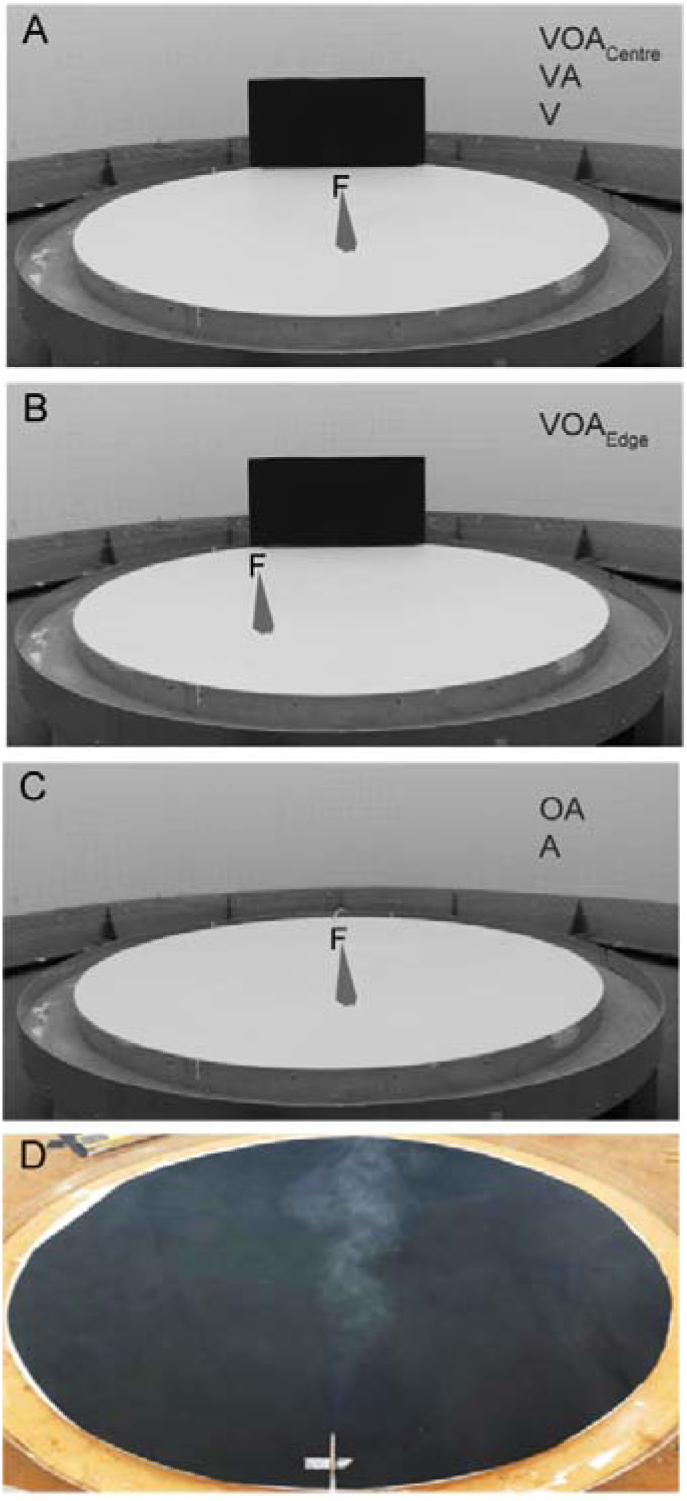
Experimental configurations. Wood ants *Formica rufa* learnt to navigate from the centre of the white platform (120 cm in diameter) to the feeder (F) 45 cm away from the centre in the presence of a visual cue (V, black rectangle), olfactory cue (O, hypothetical odour plume shown in grey) and airflow (A, airflow shown in grey) either presented individually (V or A) or together (VOA_Centre_, VOA_Edge_, VA or OA). An ant released at the centre of the platform walking towards the edge of the platform reaches the filter paper with the odour at r = 44 cm, the feeder at r = 45 cm, the centre of the visual cue together with the end of the tube connected to the pump at r = 50 cm and the edge of the platform at r = 60 cm. Each group of ants was trained to only one of the 6 different training conditions. **A:** Feeder was placed at the centre of the visual cue with either all the cues presented together (VOA_Centre_), only the visual cue and the airflow (VA) or the visual cue only (V). **B:** Feeder was placed at the left edge of the visual cue where the olfactory cue and airflow was placed (VOA_Edge_). **C:** Feeder location was defined by either the airflow only (A) or the airflow together with the odour (OA). **D:** Visualisation of the odour plume. Draeger air flow tester (tube visible at the lower end of the picture) was used to release smoking sulfuric acid and was connected to the same pump used for providing the air flow during the behavioural experiments. Platform was covered with black paper for smoke visualisation.

### Data analysis

Training and test paths were analysed from reliable ants that approached the feeder +/- 10° in at least 2/3 of their training trials from training round 14 onwards.

The performance of naïve ants experiencing the cues for the first time and from well-trained ants was determined. In the VOA experiments, tests with either the olfactory (V+O-A+) or visual cue (V-O+A+) removed were analysed also. General walking speed and path straightness (index of straightness = beeline distance / walking distance) were calculated for paths from r = 10 cm to r = 35 cm (r being distance from the centre of the platform). Accuracy of path headings were determined at r = 14 cm, 21 cm, 28 cm, 35 cm and 42 cm. To do so, the difference between the actual heading direction and the beeline to the feeder was calculated for each distance. r = 42 cm has a lower sample size because the tracking system often stopped in the close vicinity of the feeder.

A more detailed path analysis was done for all training paths from ants using different cue combinations. Here, paths were taken from the edge of the holding chamber (r = 3.25) up to r = 35 cm. We took the reliable ants (see above for selection) and focused on their paths when approaching the feeder +/- 10°. Paths were broken into 15 cm chunks and walking speed and path straightness were calculated for each of these chunks. Means for first and second halves of route were calculated for each path and means from all training paths were determined for each ant. Turns along the paths were calculated by finding 1 cm chunks with directions more than +/- 60° away from the target direction. Turns were counted for the first and second half of the route and means over all the training paths were calculated for each ant.

## Results

### Cur binding can be seen in multimodal navigation

Here we tested cue binding (as in (Steck *et al.* 2011)) in wood ants navigating to a feeder defined by a visual cue (V), olfactory cue (O) and an airflow (A) presented together (VOA training). In the first experiment, the feeder together with the odour and airflow was placed in the centre of the visual cue (see VOA_Centre_ in Fig. 1A). To accurately approach the feeder location during multimodal navigation, experienced wood ants required all the individual parts of the learnt cue combination. VOA ants that were trained to find a feeder defined by a visual cue (V), olfactory cue (O), and airflow (A) together were subsequently tested with OA or VA only. With either the visual (V) or olfactory (O) cue removed in these tests, well-trained ants were not able to approach the feeder accurately (V+O-A+ and V-O+A+ in Fig. 2). Hence, the cues were bound together, and individual parts of the learnt combination alone were not anymore sufficient to locate the familiar feeder location. The observed decrease in accuracy (Fig. 2D) was coupled with a lower path straightness (Fig. 2F) and walking speed (Fig. 2E). Importantly, both path straightness and walking speed were lower than in ants that were trained with OA or VA only (Mann Whitney U tests; Index of straightness, V-O+A+ vs OA, p < 0.001, V+O-A+ vs VA, p < 0.001; walking speed, V-O+A+ vs OA, p < 0.05, V+O-A+ vs VA, p < 0.001).

**Figure 2:**
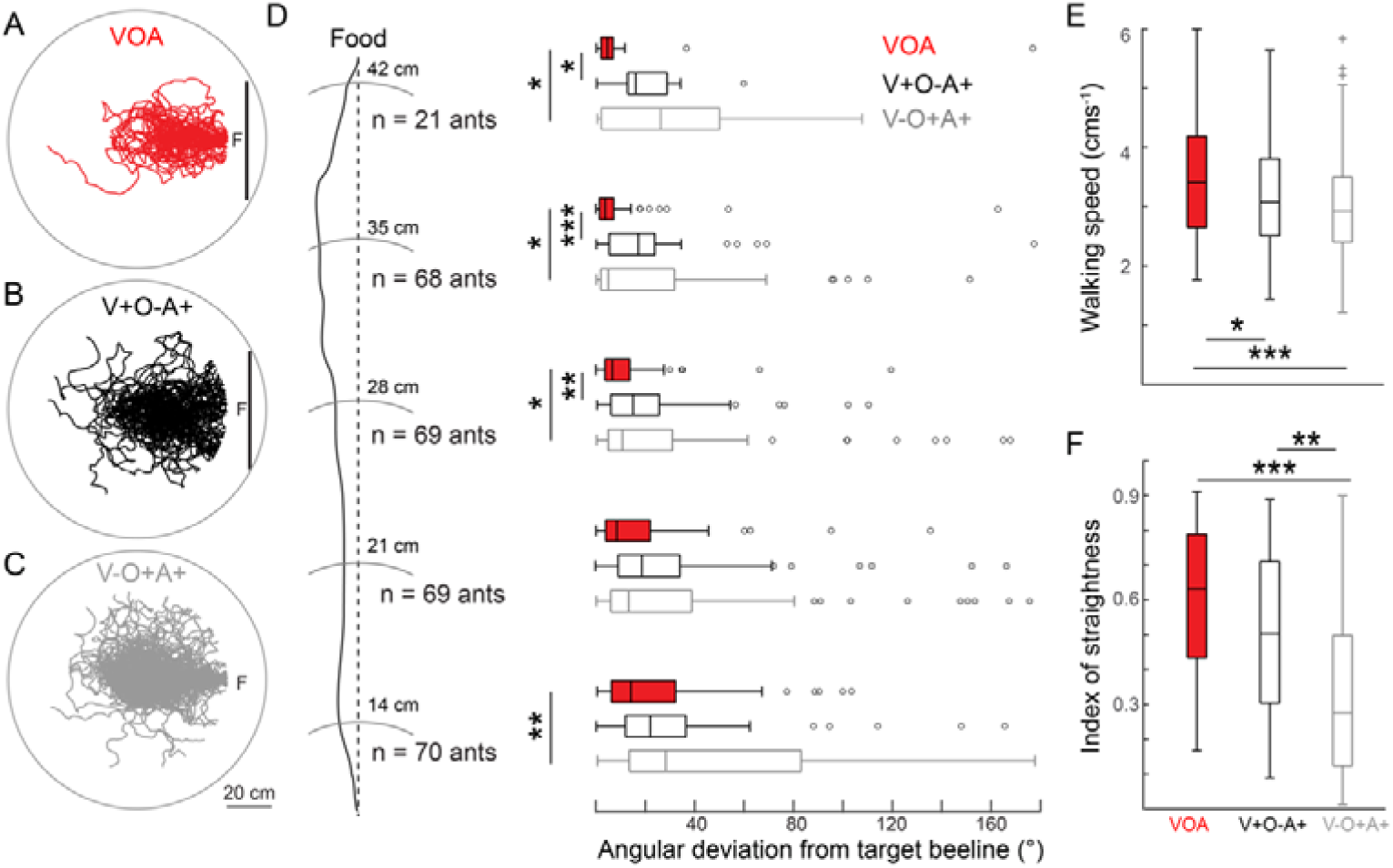
All individual learnt cues are needed for accurate multimodal navigation. **A:** Ants learnt to navigate to a feeder (F) that was 45 cm away from the centre of the platform (120 cm in diameter) and defined by a visual cue (V, shown as a black bar), olfactory cue (O) and an airflow (A). The feeder was placed at the centre of the visual cue (VOA_Centre_). Trajectories from individual ants are shown as red lines. **B:** Ants trained with VOA were tested with the odour removed (V+O-A+). Trajectories from individual ants are shown as black lines. **C:** Ants trained with VOA were tested with the visual cue removed (V-O+A+). Trajectories from individual ants are shown as grey lines. **D:** Path accuracy was calculated along the paths at r = 14 cm, 21 cm, 28 cm, 35 cm and 42 cm. For each distance, the ants’ angular deviation from the beeline to the feeder (shown as dashed line) are shown as boxplots (median, 25th and 75th percentiles (edges of the boxes) and whiskers for extreme values not considered as outliers (°)). The p values from a Friedman test with Dunn’s multiple comparison tests are shown as asterisks for each distance (*, p < 0.05; **, p < 0.01; ***, p < 0.001). For any given distance, ants are only considered if they provide a value for all 3 conditions. **E:** Walking speed for training (VOA) and the two test conditions (V+O-A+ and V-O+A+). n = 69 ants. p values for a Friedman test with Dunn’s multiple comparison tests are shown as asterisks. For boxplot and p value conventions see **D. F:** Path straightness for training and tests. Index of straightness values are between 0 and 1, with 1 indicating a perfectly straight path. n = 69 ants. For boxplot and p value conventions see **D**.

### Cue binding depends on the navigational context

We further showed that the observed cue binding is not a rigid property of multimodal cue learning and depends on the navigational context. Another group of ants was trained with the same set of cues (OVA), but the feeder together with the odour and the airflow was now placed at the edge of the visual cue (Fig. 1B) to create a simpler navigational task (Harris *et al.* 2007). In contrast to what we have seen before (see Fig. 2), well-trained ants were still able to accurately approach the learnt feeder location when either the olfactory (O) or visual (V) cue was removed (V+O-A+ and V-O+A+ in Fig. 3). However, when far away, ants were less accurate with the visual cue removed (Fig. 3D). Furthermore, path straightness was decreased (Fig. 3F) but walking speed was not altered (Fig. 3E).

**Figure 3:**
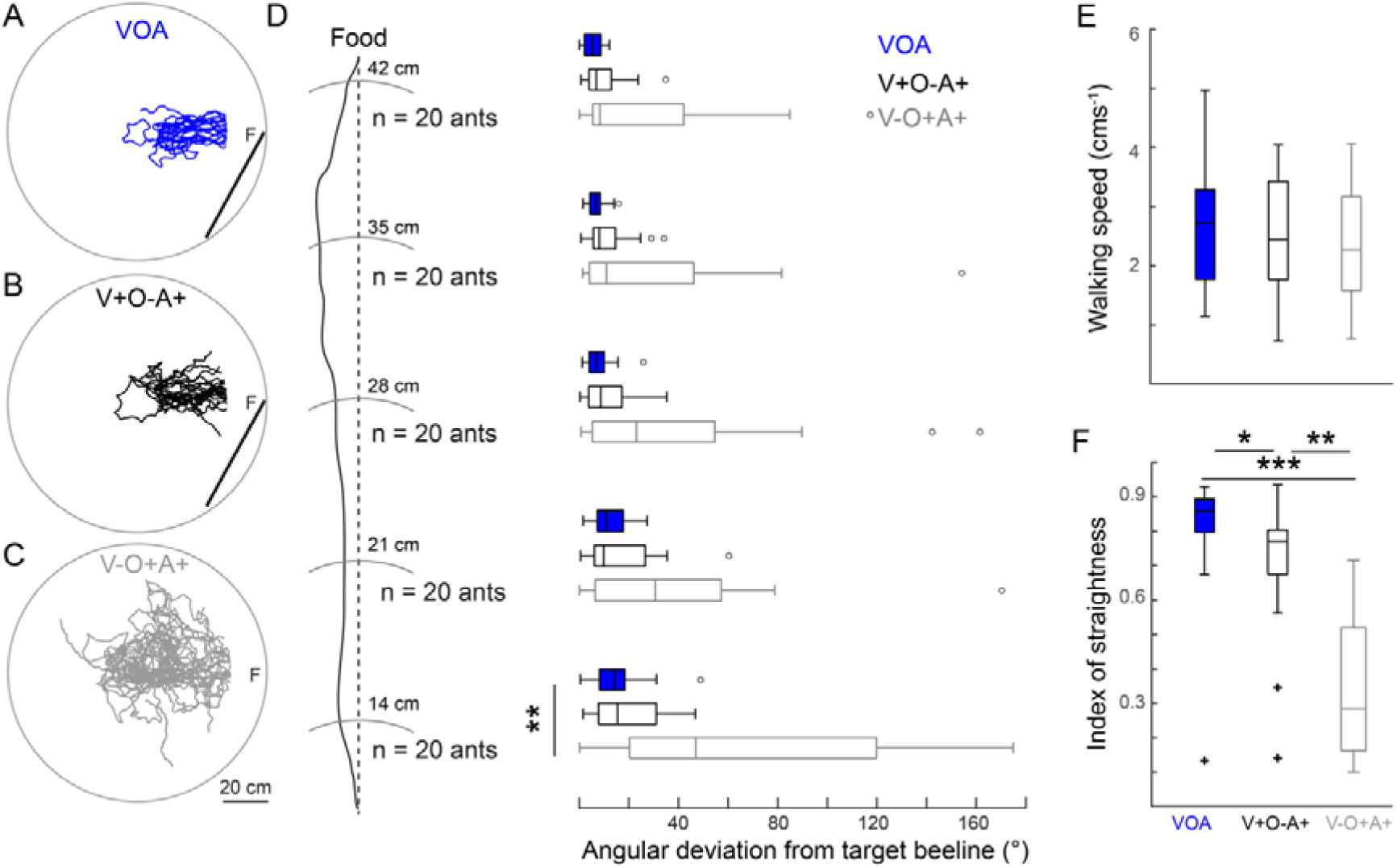
Multimodal cue binding is flexible. **A:** Ants learnt to navigate to a feeder (F) that was 45 cm away from the centre of the platform (120 cm in diameter) and defined by a visual cue (V, shown as a black bar), olfactory cue (O) and an airflow (A). The feeder was placed at the left edge of the visual cue (VOA_Edge_). Trajectories from individual ants are shown as red lines. **B:** Ants trained with VOA were tested with the odour removed (V+O-A+). Trajectories from individual ants are shown as black lines. **C:** Ants trained with VOA were tested with the visual cue removed (V-O+A+). Trajectories from individual ants are shown as grey lines. **D:** Path accuracy was calculated along the paths at r = 14 cm, 21 cm, 28 cm, 35 cm and 42 cm. For each distance, the ants’ angular deviation from the beeline to the feeder (shown as dashed line) are shown as boxplots (median, 25th and 75th percentiles (edges of the boxes) and whiskers for extreme values not considered as outliers (°)). The p values from a Friedman test with Dunn’s multiple comparison tests are shown as asterisks for each distance (*, p < 0.05; **, p < 0.01; ***, p < 0.001). For any given distance, ants are only considered if they provide a value for all 3 conditions. **E:** Walking speed for training (VOA) and the two test conditions (V+O-A+ and V-O+A+). n = 20 ants. p values for a Friedman test with Dunn’s multiple comparison tests are shown as asterisks. For boxplot and p value conventions see **D. F:** Path straightness for training and the tests. Index of straightness values are between 0 and 1, with 1 indicating a perfectly straight path. n = 20 ants. For boxplot and p value conventions see **D**.

### Cues act additively in determining paths accuracy after training

Across the different experimental conditions, we saw some innate attraction to the cues in naïve ants (Fig. 4). In all conditions with the visual cue present, ants were directed towards it (Rayleigh test, all p < ***) whilst the odour or the airflow were not attractive to them (Rayleigh test, both p > 0.05). Training improved the ants’ navigational performance, and in general, experienced ants walked more accurately towards the feeder (Angular deviation from target beeline at r = 35 cm; Wilcoxon matched pairs test, VOA_C_ p < 0.001, VOA_E_ p < 0.001, VA p < 0.001, OA p < 0.001, V p < 0.001), had straighter paths (Index of straightness; Wilcoxon matched pairs test, VOA_C_ p > 0.05, VOA_E_ p > 0.05,VA p < 0.001, OA p < 0.001, V p < 0.001) and a lower walking speed (Walking speed; Wilcoxon matched pairs test, VOA_E_ p < 0.01, VOA_C_ p < 0.01, VA p < 0.001, OA p < 0.001, V p < 0.05) than naïve ants unfamiliar with the experimental environment (for paths and heading direction see Fig. 4). Comparing the different cue combinations, we see that the proportion of accurate ants was increased with the number of navigational cues available (Fig. 5A), i.e. cues act additively in determining paths accuracy.

**Figure 4:**
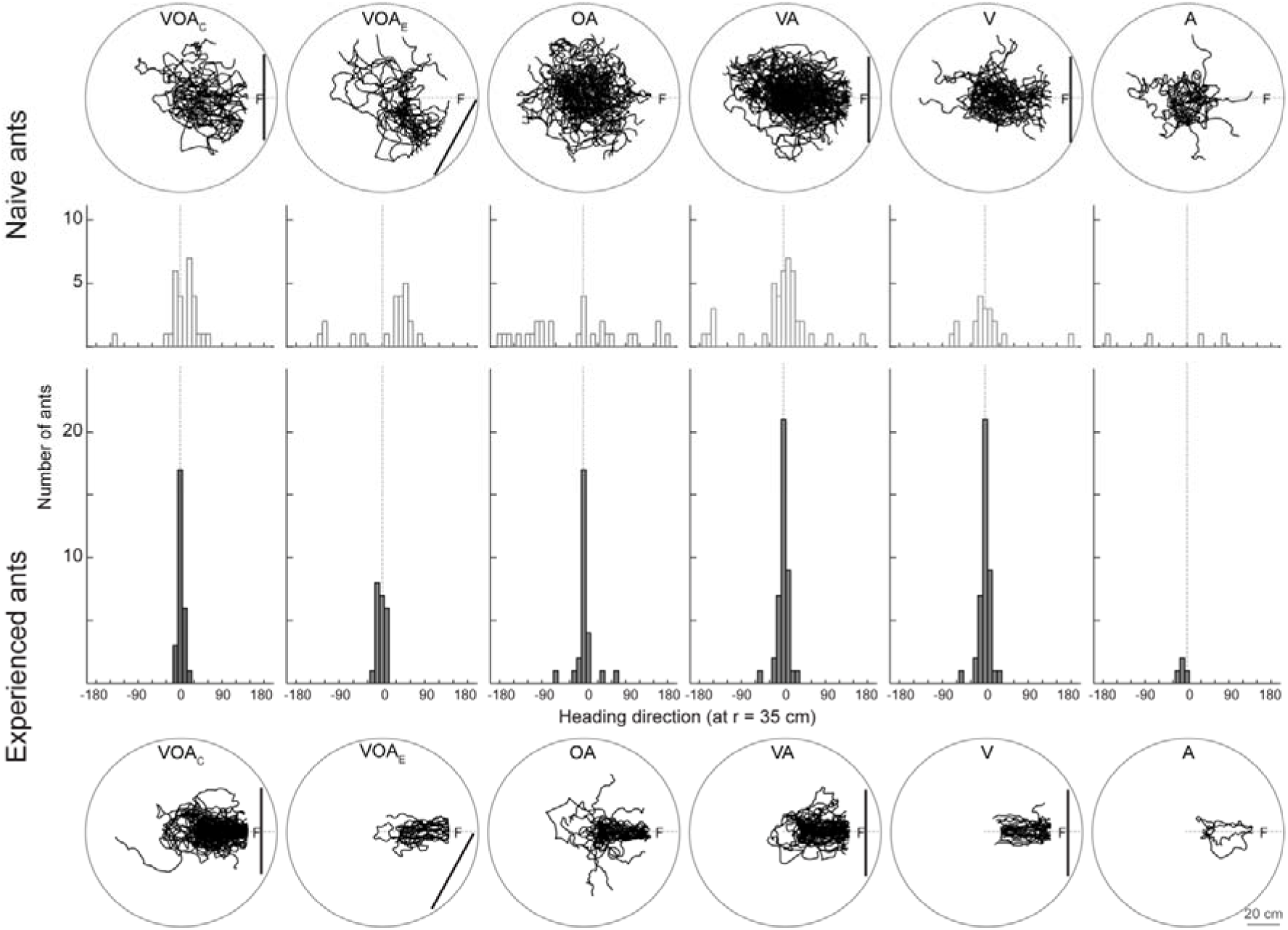
From innate attraction to stable routes. Naïve ants experiencing the cues for the first time (top, VOA_Centre_: n = 27 ants, VOA_Edge_: n = 24 ants, OA: n = 27 ants, VA: n = 44 ants, V: n = 20 ants, A: n = 5 ants) and well-trained ants after an extensive training (bottom, VOA_Centre_: n = 88 ants, VOA_Edge_: n = 22 ants, OA: n = 32 ants, VA: n = 53 ants, V: n = 22 ants, A: n = 4 ants) are shown for each training condition. Trajectories from individual ants are shown as black lines. The feeder (F) was 45 cm away from the centre of the platform (120 cm in diameter) and defined by a visual cue (V, shown as a black bar), olfactory cue (O) and an airflow (A) presented individually of together. Each group of ants was only trained in one of the 6 conditions. The heading direction at r = 35 cm from ants that were recorded in both conditions are shown in the histogram in the middle (bin size: 10°). Because naïve ants were not recorded for all experimental groups, the sample size for the statistics is lower than in the figure to allow pairwise comparisons. VOA_Centre_: n = 27 ants, VOA_Edge_: n = 22 ants, OA: n = 27 ants, VA: n = 42 ants, V: n = 19 ants, A: n = 4 ants. For statistics see results.

**Figure 5:**
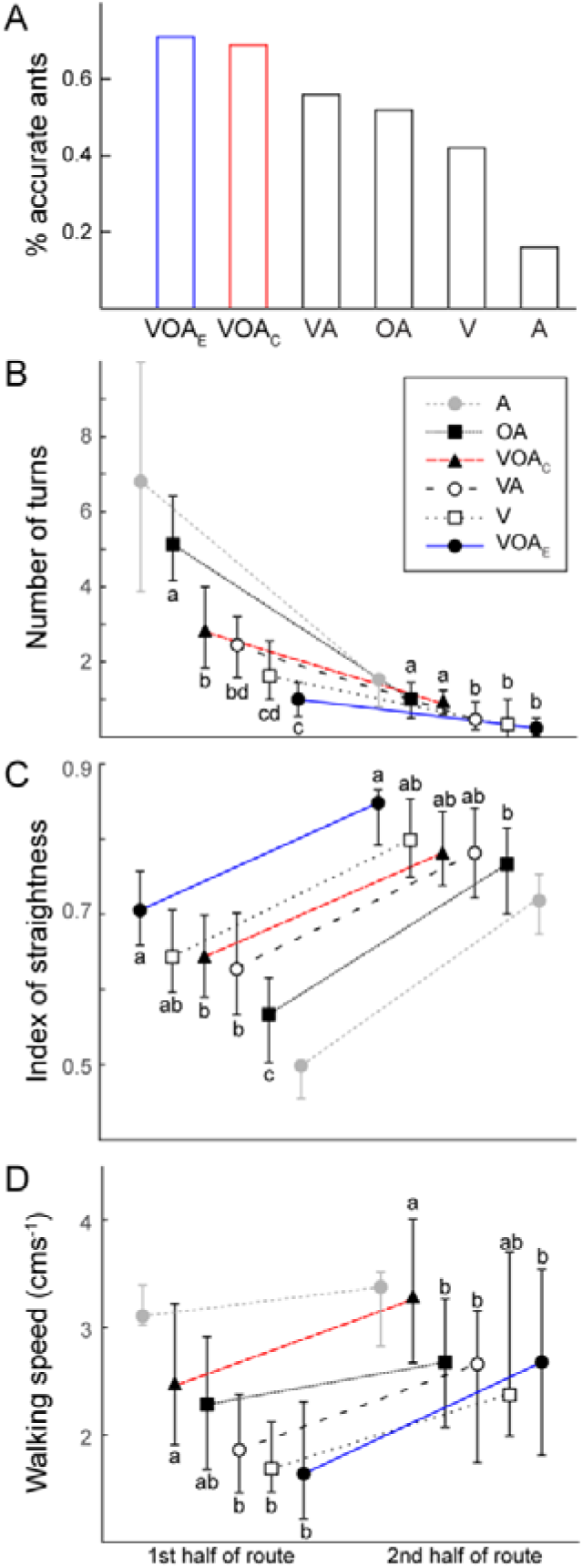
Path characteristics of ants using different cues for navigation. **A**: Proportion of accurate ants for the six different experimental conditions. Accuracy is defined as the proportion of ants that approached the feeder +/- 10° in at least 2/3 of their training trials. VOA_Edge_: 24 out of 34 ants, VOA_Centre_: 88 out of 128 ants, VA: 56 out of 100 ants, OA: 32 out of 62 ants, V: 25 out of 60 ants, A: 5 out of 32 ants. **B:** Number of turns shown for the different training conditions for the first and second part of the route to the feeder. Data are plotted as medians with error bars showing the 25th and 75th percentiles. For different training conditions see inset. Lowercase letters indicate significant differences (p<0.05) between the different cue combinations (Kruskal Wallis with Dunn’s multiple comparison tests). Groups with same letters are not significantly different. Ants trained with the airflow (A) are not included in statistics because of low sample size. VOA_Edge_: n = 24 ants, VOA_Centre_: n = 88 ants, VA: n = 56 ants, OA: n = 32 ants, V: n = 25 ants, A: n = 5 ants. **C:** The same as **B** but for path straightness. **D:** The same as **B** but for walking speed.

### Different navigational strategies require different path structures

Further path analysis of well-trained ants revealed that the learning of different cues produces differently structured paths. Generally, ants had fewer turns (Fig. 5B), straighter paths (Fig. 5C) and a higher walking speed (Fig. 5D) in the second half of their route to the feeder (Wilcoxon matched-pairs signed-ranked test, all p < 0.001). When comparing the different conditions, we observed that ants had more sinuous paths and produced more turns when they followed the odour plume but did not have any visual information available (Fig. 5B and 5C). Contrary to this, V and VOA_Edge_ ants had the straightest paths and the lowest number of turns (see Fig. 5B and 5C). Interestingly, ants with exactly the same set of navigational cues (VOA), but having the feeder either in the centre (VOA_Centre_) or at the edge (VOA_Edge_) of the visual cue, differed from each other. The VOA_Edge_ ants had straight paths and only a few turns, whereas the VOA_Centre_ ants had paths somewhere in between visually and olfactory guided ants. Moreover, VOA_Centre_ ants walked significantly faster than most of the other groups while VOA_Edge_ ants were in the lower speed range together with the other groups (Fig. 5D).

### Functional range of the odour plume is bigger than of the airflow

In an additional test, ants from the VOA_Centre_ training were tested with the odour and airflow (OA) shifted 20° to the left (Fig. 6A and B for the two different conditions). These ants initially headed towards the centre of the visual cue (see orange arrow in Fig. 6A and B) and then drifted away from this direction to walk towards the shifted OA (see blue arrow in Fig. 6A and B). Hence, visual information was used for navigation when far away and the odour plume became more important further along the route. Ants from the VA training were tested in a similar conflict test, i.e. with airflow (A) displaced by 20°, and they also initially walked towards the centre of the visual cue and then shifted away towards the source of the airflow (Fig. 6C). However, the change of direction happened later along the route suggesting that the spatial scale over which the airflow can be detected, and used as a cue, is smaller than the one for the odour plume.

**Figure 6:**
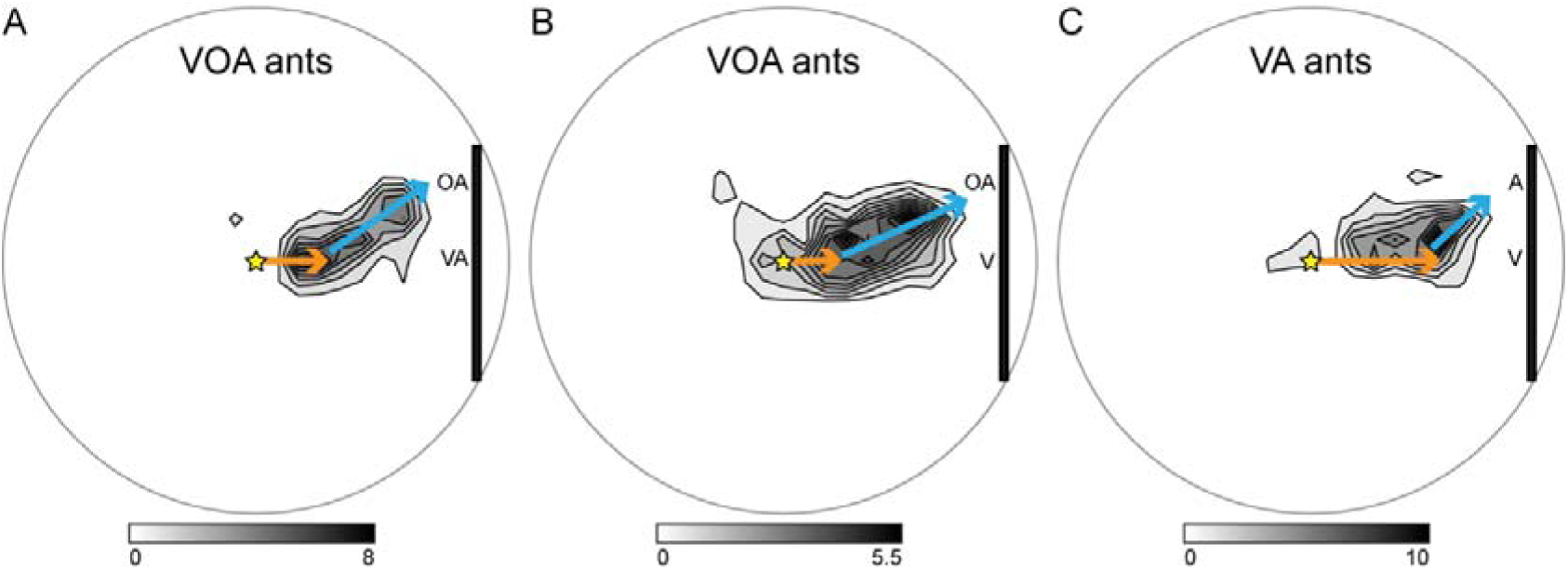
Routes taken by trained ants in cue conflict tests. Path density plots are shown for three different groups of ants. **A**: VOA_Centre_ ants were tested with the olfactory cue and airflow (OA) shifted 20° to the left and a clean airflow (A) added to the centre of the visual cue (VA). **B**: VOA_Centre_ ants were tested with OA shifted 20° to the left and only the visual information in the centre (V). **C**: VA ants were tested with the airflow (A) shifted 20° to the left. Asterisk: point of release. Black rectangle: visual cue. Orange arrow: intial walking direction. Blue arrow: change of walking direction.

## Discussion

Ants are excellent multimodal navigators (Knaden & Graham 2016) and here we present information about the details of multimodal cue use in navigating wood ants *Formica rufa*. Controlled lab experiments with detailed recording of ants’ movements have allowed us to investigate a series of questions about the use of unimodal or multimodal cues to find a feeder. We show that (i) the navigation performance is improved when multiple cues are available, (ii) different navigational strategies require different movement patterns and (iii) the cue binding previously described by (Steck *et al.* 2011) is flexible and context dependant, with the nature of cue binding potentially being explained by the sensori-motor contingencies of a particular task.

### Multimodal sensori-motor integration

Wood ants are excellent navigators, foraging for dead arthropods on the ground and aphid honeydew in trees (Domisch *et al.* 2016). We know that vision is important for navigation in many ants (Zeil 2012; Wehner *et al.* 2014), including wood ants (Rosengren 1971) but for the detection of food sources an excellent sense of smell is also essential (dead arthropods: (Buehlmann *et al.* 2014); aphids: (Nault et al. 1976; Lohman et al. 2006; Verheggen et al. 2012)). Here, we show that ants utilise these ecologically relevant sensory modalities, to more accurately approach a learnt feeder when multiple cues were available (Fig. 5A) and in general we see cues acting additively in determining the ants’ path accuracy. This demonstration of more efficient paths when multiple cues are available, adds to the weight of evidence on the value of multimodal information in a range of behaviours. Multimodal integration has already been shown to enhance performance in perception (van Swinderen & Greenspan 2003; Goyret *et al.* 2007; Chow & Frye 2008; van Breugel & Dickinson 2014) and learning (Rowe 2002; Guo & Guo 2005; Reinhard *et al.* 2006; Steck *et al.* 2011) in insects.

We know from fruit flies and other flying insects that visual feedback is needed for stabilizing an upwind flight (Reiser *et al.* 2004; Budick *et al.* 2007), thus plume tracking is enhanced in the presence of visual cues (Fadamiro *et al.* 1998; Frye *et al.* 2003) where the cross modal interaction works because attractive odours enhance the gain of optomotor responses during flight (Chow & Frye 2008) and with more precise flight, it is easier to track spatial odour gradients (Duistermars & Frye 2010; Stewart *et al.* 2010). In a potential similarity, we observed that the use of different cues impacts on the ants’ movement patterns (Fig. 5B, C and D). Across all conditions, experienced ants that accurately approached the learnt feeder and always walked slower and straighter than naïve ants on their first time experiencing the cues (Fig. 4). This suggests that a lower walking speed is useful for accurate navigation. Indeed, we have seen in a previous study with *C. fortis* desert ants that low walking speeds correlate with the learning and use of sensory information (Buehlmann et al. 2018). More detailed path analysis further revealed that navigating wood ants had more sinuous paths with more turns when they followed the odour plume but did not have any visual information available (see OA in Fig. 5B and 5C) and adding a visual cue to the odour information positively impacted on plume following (see OA vs VOA_Centre_) similarly to flies (Duistermars & Frye 2010; Stewart *et al.* 2010).

Further similarities with flying insects can be highlighted from the conflict tests, where odour and airflow were displaced relative to the visual cue (Fig. 6). When airflow alone was displaced (Fig. 6C) the change of direction happened later along the route than for airflow and odour (Fig. 6A and B), suggesting that the spatial scale over which the airflow can be detected is smaller than the one for the odour plume. We have learnt from studies in mosquitoes how cues can integrate in a sequential manner. Mosquitoes have developed an elegant mechanism to respond to the multiple cues that indicate a host. They sense carbon dioxide from far away and this activates a strong attraction to visual cues which allows mosquitoes to approach a host and then when closer to the target they eventually approach it using thermal cues (van Breugel et al. 2015). Hence, different spatial scales for the different cues allows the insects to use the cues sequentially while at the same time cues interact

### Is cue binding a cognitive process or can it be explained by sensory motor motifs?

In the experiments presented here, we challenged ants to learn a feeder location defined by multimodal information. Previously, in a similar experiment, desert ants learnt to use both olfactory and visual cues to navigate back to their nest (Steck *et al.* 2011) and the bimodal cues were first learnt independently but later stored as a unit, i.e. the components were merged and ants no longer responded to either visual or olfactory cue presented alone (Steck *et al.* 2011). We ask here if this cue binding can be explained by the ants’ path characteristics and if some behavioural flexibility is retained.

We performed experiments where the feeder location was defined by a visual cue (V), olfactory cue (O) and airflow (A) presented together (VOA experiments). In the first experiment, the feeder was located in the centre of the visual cue (VOA_Center_). These ants were not anymore able to accurately approach the feeder when either the visual component or the olfactory component was removed, they also walked slower and less straight (Fig. 2). Hence, as previously described (Steck *et al.* 2011), all the learnt cues were required for accurate navigation. However, in a second experiment ants had the same set of cues (VOA) but the feeder was now located at the edge of the visual cue (VOA_Edge_). Here, ants in tests with either the olfactory or visual cue missing were not significantly less accurate (Fig. 3). Interestingly, ants from the two groups (VOA_Centre_ and VOA_Edge_) were significantly different in path straightness, turn frequency and walking speed, even though they experienced the same set of cues (Fig. 5).

In summary, we can conclude that ‘binding’ is not a cognitive inevitability in multimodal tasks, but depends on the sensori-motor contingencies of the particular task. Furthermore, the interactions between cues may be either: Direct, for instance when sensori-motor patterns in experienced ants when the removal of a familiar cue is deleterious on path efficiency; Or, indirect, where the fine-details of the sensori-motor patterns during learning are important for the ultimate performance of well-trained ants. Navigating to the edge of a large visual cue is a simpler task than navigating to the centre (Harris *et al.* 2007), thus the rapid visual learning could also increase the efficiency of olfactory learning.

## Acknowledgements

We thank Antoine Wystrach for providing the Matlab script for the path density plots and Tom Collett for many fruitful discussions.

## Competing interests

No competing interests declared.

## Funding

This project was funded by the people programme (Marie Curie Actions) of the European Union’s Seventh Framework Programme (FP7/2007-2013, under REA grant agreement no. PIEF-GA-2013-624765) and a fellowship from the Swiss National Science Foundation (grant no. P2SKP3-148476) to CB. PG and CB are additionally funded by a BBSRC grant BB/R005036/1.

